# 15 second assessment of hand and foot motor skill using a smartphone

**DOI:** 10.1101/2024.10.29.621007

**Authors:** Atsushi Takagi, Noriyuki Tabuchi, Wakana Ishido, Chikako Kamimukai, Hiroaki Gomi

## Abstract

Motor skills are essential for daily functioning and serve as key indicators of healthy development and aging. However, existing assessments of motor skill are inaccessible to the public and do not directly measure the quality of the movement. Here, we developed a convenient and robust measure of motor skill that uses a smartphone app to provide on-the-spot assessment in just 15 seconds. To validate our method, we asked 1675 participants between the ages of three to eighty-eight years old to trace circles at a fixed rhythm with a smartphone, which was either held in the hand or strapped to the ankle. Motor skill was quantified by the smartphone app using an algorithm that calculated the variability in the acceleration’s trajectory. Our assessment revealed significant changes in the skill of the hands and feet with age and practice. The variability of the hands and feet linearly decreased and matured in the mid-teens, but it regressed gradually thereafter. Laterality, or the difference in the motor skill between the left and right limbs, increased with age as the non-dominant hand and foot regressed faster in the elderly. Motor practice affected both skill and laterality as left-handers who were forced to write with their right-hand during childhood had a tell-tale sign of stronger right-handedness and, surprisingly, right-footedness. Our assessment aims to democratize motor skill assessment, making it accessible for professionals and users of all ages.

## Introduction

Our daily lives and wellbeing are dependent on the ability to move around and interact with the environment. Tasks of daily living such as writing and walking demand the skillful use of the hands and feet. Existing technologies have measured the motor skill of the hands and feet by documenting a participant’s performance at a select motor task. An example is the peg hole test where participants insert as many small pins into thin holes as possible in thirty seconds^1,2^. Another assessment records the number of blocks moved from one box to another in a given duration^3^. The maximum number of index finger taps in a given time span has also been used to gauge a hand’s skill^4^. Another measures the maximum number of circles filled with a pencil^5^. Fewer tests are available for the lower-limb, but some examples include the maximum number of foot taps^6^ and the ability to balance stably on one leg^7^.

These assessments of motor skill can serve as important markers of development in children and of decline in the elderly. Studies have shown how the performance in inserting pegs with the hands improves with growth^8^ and regresses with aging^2^. Gasser et al. performed a battery of motor tests like the maximum repetitive and alternating movement of hands and feet, the pegboard test, and static and dynamic balance with children. They showed that the performance of most tests improved and plateaued at adolescence^9^.

Motor skill assessments are important for parents, guardians, and health professionals in tracking the development of a child’s motor skill and gauging the decline in an aging individual’s skill when moving the hands and feet. However, existing assessments of hand and foot skill require some combination of dedicated equipment and a trained professional to administer and assess the outcome of the test, which takes time and is costly. Furthermore, existing methodologies only provide an indirect measure of motor skill. The quality of a limb’s movement can be characterized by its accuracy or precision, which is dependent on the ability to repeat a movement without variability^10,11^. What is needed is an assessment of motor skill that does not require a trained professional to administer, uses equipment that is widely available, and one that measures the quality of the limb’s movement.

The goal of this study was to develop an accessible assessment of motor skill based on the movement of the hands and feet. We opted to use a smartphone device as it comes equipped with an inertial measurement sensor and is used by approximately 4 billion people or half of the global population^12^. Since a change in movement variability at the same speed signifies an improvement in skill^10,13,14^, we measured motor skill by asking participants to trace a circle using their hands or feet at a constant rhythm set by a metronome whilst a smartphone was held in their hand or strapped to their ankle (Figure 1a). Preliminary tests were conducted with children and the elderly to find speeds for the hands and the feet that all participants felt comfortable with, which was 2.5 Hz for the hands and 1.5 Hz for the feet. The circles they had to trace had a diameter of 10 cm and 15 cm for the hands and feet, respectively. A smaller smartphone was used for kids aged 12 and under so that it could be held comfortably in the hand. The smartphone’s three-axis accelerometer recorded each limb’s acceleration trajectory. We developed a novel algorithm that calculated the variability of the acceleration trajectory within the smartphone (see Methods).

**Figure 1.**
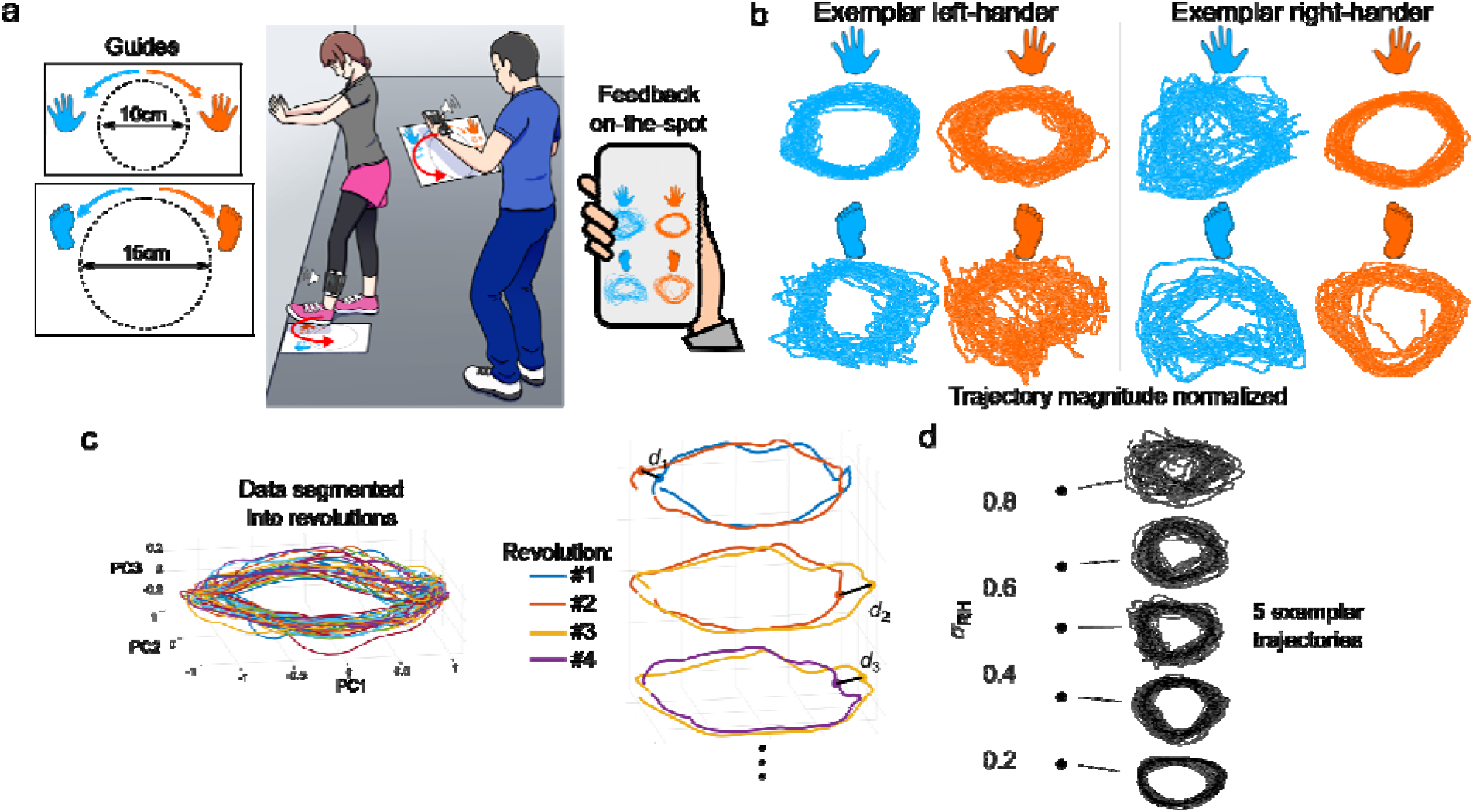
Motor skill of the hands and feet was quantified by acceleration variability during periodic circle tracing. (a) Smartphone was either held or strapped to the ankle. Participants traced a circle in the outwards direction in time with a metronome for fifteen seconds at 2.5 Hz for the hands and at 1.5 Hz for the feet. Feedback of motor skill can be provided on-the-spot by the same device. (b) Hand and foot acceleration trajectories from an exemplar left-hander and right-hander. The normalized trajectory was projected onto the 2D surface consisting of its two major principal vectors (actual trajectory is 3D). (c) Procedure to calculate acceleration variability. PCA was used to reorient the acceleration trajectory along the dimensions of greatest variance, then the trajectory was segmented into revolutions for each circular motion. Hausdorff distance *d* was calculated between consecutive revolutions, and the average Hausdorff distance of all comparisons yielded the acceleration variability. (d) Right-hand’s acceleration variability and corresponding trajectory from five participants. Acceleration variability increases as the trajectory’s variability visibly increased.

## Results

1675 participants took part in the study with ages ranging from three to eighty-eight years old. Participants moved the smartphone to trace circles with either their hands or their feet in time with the smartphone’s metronome (Figure 1a, see Methods for details). Once movement data was collected from both arms and legs, participants completed a questionnaire that contained the Edinburgh Handedness Inventory^15^ and the Coren Footedness Inventory^16^. They were also asked whether they were forced to write with their right-hand as a child.

The acceleration along the three-dimensional axes was recorded throughout the circular movement. The acceleration trajectory’s offset was removed to centralize the trajectory, then its size was normalized by dividing the values by the mean Euclidean norm. Principal component analysis was then used to orient the trajectory along its first two principal components (see Methods). Upon visual inspection, the acceleration trajectory from a strongly left-handed (Edinburgh quotient of -1) and a strongly right-handed person (Edinburgh quotient of +1) suggested greater variability in the non-dominant limb’s acceleration trajectory (Figure 1b). To avoid the effects of drifts in the acceleration sensor, the limb’s variability was quantified by segmenting the acceleration trajectory into revolutions, where one revolution corresponded to one period of the circular movement (Figure 1c, left). The start and end of a revolution were when the acceleration vector’s angle crossed the axis of the first principal component PC1 (Figure 1c). From each fifteen second recording, approximately thirty-four revolutions from the hand and twenty revolutions from the feet were identified. Since the sampling rate of the acceleration sensor was variable, we calculated the Hausdorff distance between consecutive revolutions, a spatial measure of distance commonly used to identify the similarity of images^17^ (Figure 1c, right). Roughly thirty-three and nineteen Hausdorff distances were calculated from the hand and foot recordings. The average Hausdorff distance yielded the movement variability of the left hand *σ*_LH_, right hand *σ*_RH_, left foot *σ*_LF_, and right foot *σ*_RF_ . Figure 1d shows the right-hand’s acceleration trajectory from five participants with varying degrees of acceleration variability ranging from *σ*_RH_ = 0.2 to 0.8, where smaller variability corresponds to greater motor skill. The acceleration trajectory becomes visibly more variable as *σ*_RH_ increases.

To quantify the limb’s variability at the same speed, we restricted all further analysis to participants who completed the circular movements within ±150 milliseconds of the target period, leaving n=1413 data for the hands and n=1196 for the feet. We then examined how the acceleration variability of the non-dominant and dominant limbs changed with age. The dominant and non-dominant hands were identified using the Edinburgh quotient^15^. An Edinburgh quotient smaller than zero meant that the left-hand was the dominant hand and the right-hand was the non-dominant one, and vice versa when the Edinburgh quotient was greater than zero. A similar process was used to identify the dominant and non-dominant feet using the Coren quotient^16^.

The acceleration variability was binned and averaged across a span of two neighboring years, yielding one point for every two years (Figures 2a and 2b, error bars signify mean and one standard error). The acceleration variability of all limbs decreased linearly and rapidly from age four until the age of maturation *Y*_M_, which was a critical point as the decrease in acceleration variability stopped abruptly. We estimated the age *Y*_M_ when the acceleration variability matured by linearly fitting the data until the age of maturation *Y*_M_. A second-order polynomial was used to fit the data above *Y*_M_ because the second-order term significantly improved the fit for both the hand and foot acceleration variability, but a third-order term did not (see Methods for details). We identified the value of *Y*_M_ that minimized the least-squares distance between the fitted acceleration variability and the data for the non-dominant and dominant limbs together. This process was carried out separately for the hands and the feet. Bootstrap resampling method with 10000 samples was used to estimate the 95% confidence interval of *Y*_M_^18^ (see Methods). The skill of the hands matured at *Y*_M_ = 13 ± 1.4 and the skill of the feet matured at *Y*_M_ = 15 ± 0.7 (mean±95% confidence interval of bootstrapped means).

**Figure 2.**
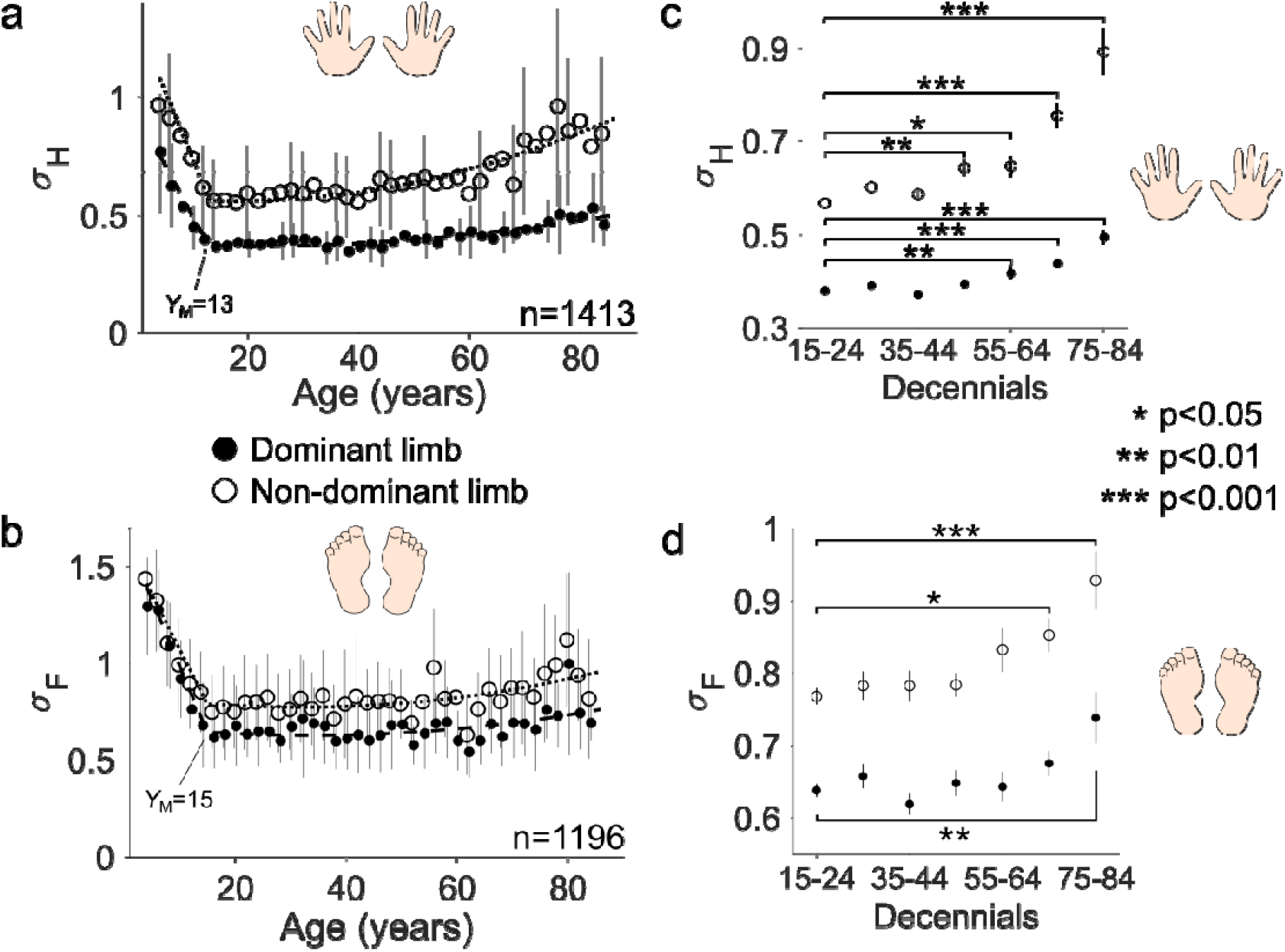
Hand and foot acceleration variability decreases with in children and regresses with aging. (a) Hand acceleration variability as a function of age (each point is the mean and standard error from two-year bins). Acceleration variability declined and matured at, then gradually regressed with aging. Dashed lines are a linear fit of the data until and a second order fit thereafter. (b) Foot acceleration variability followed a similar pattern to hand acceleration variability. (c) Hand acceleration variability separated into decennials or ten-year bins. Non-dominant hand’s acceleration variability regressed faster relative to the dominant hand when comparing with the youngest decennial. (d) Foot acceleration variability as a function of decennials. Acceleration variability of the non-dominant foot regressed faster than the dominant foot.

To estimate the age at which the acceleration variability began to regress, we grouped the data into decennials aged 15–24, 25–34, 35–44, 45–54, 55–64, 65–74 and 75–84 years old. Four one-way analyses of variance were carried out separately for each limb with the decennial as the predictor. The analysis revealed a significant effect of the decennial group on the acceleration variability of the dominant hand (F(6,990)=24.2, p<0.001), non-dominant hand (F(6,990)=29.7, p<0.001), dominant foot (F(6,886)=3.5, p=0.002), and non-dominant foot (F(6,886)=5.7, p<0.001). To see where the acceleration variability began to regress significantly, we carried out post-hoc tests between the youngest decennial group (15–24 years old) against all other decennial groups. The post-hoc tests were controlled for multiple comparisons using Tukey’s HSD. The non-dominant hand regressed the fastest as a significant difference was observed between the youngest decennial group and the 45–54-year-old decennials (Figure 2c). The dominant hand regressed slower as a significant difference was only observed after 55–64-year-old decennials. Similarly, the non-dominant foot regressed faster than the dominant foot (Figure 2d). To summarize, both the non-dominant hand and foot regressed faster than the dominant limbs.

A clear difference in the variability between the left and right limbs was observed in both the hands and the feet. Was this difference related to the handedness and footedness of each participant? We calculated the difference in the acceleration variability between the left and right hands *σ*_LH_ − *σ*_RH_ and for the feet *σ*_LF_ − *σ*_RF_, then plotted them as a function of the Edinburgh quotient^15^ and the Coren quotient^16^ (Figures 3a and 3b). Both quotients span between -1, implying strong left-handedness or footedness, to +1, which indicates strong right-handedness or footedness. We fit two linear-mixed effects model with the difference in acceleration variability as the dependent variable and the quotient as the predictor to examine the relationship between them. A significant positive linear relationship was observed between *σ*_LH_ − *σ*_RH_ and the Edinburgh quotient (Figure 3a, slope=0.21±0.01, p<0.001), and between *σ*_LF_ − *σ*_RF_ and the Coren quotient (Figure 3b, slope=0.05±0.01, p<0.001). Thus, the hand or foot mainly used in tasks of daily living had a lower acceleration variability relative to the other limb.

**Figure 3.**
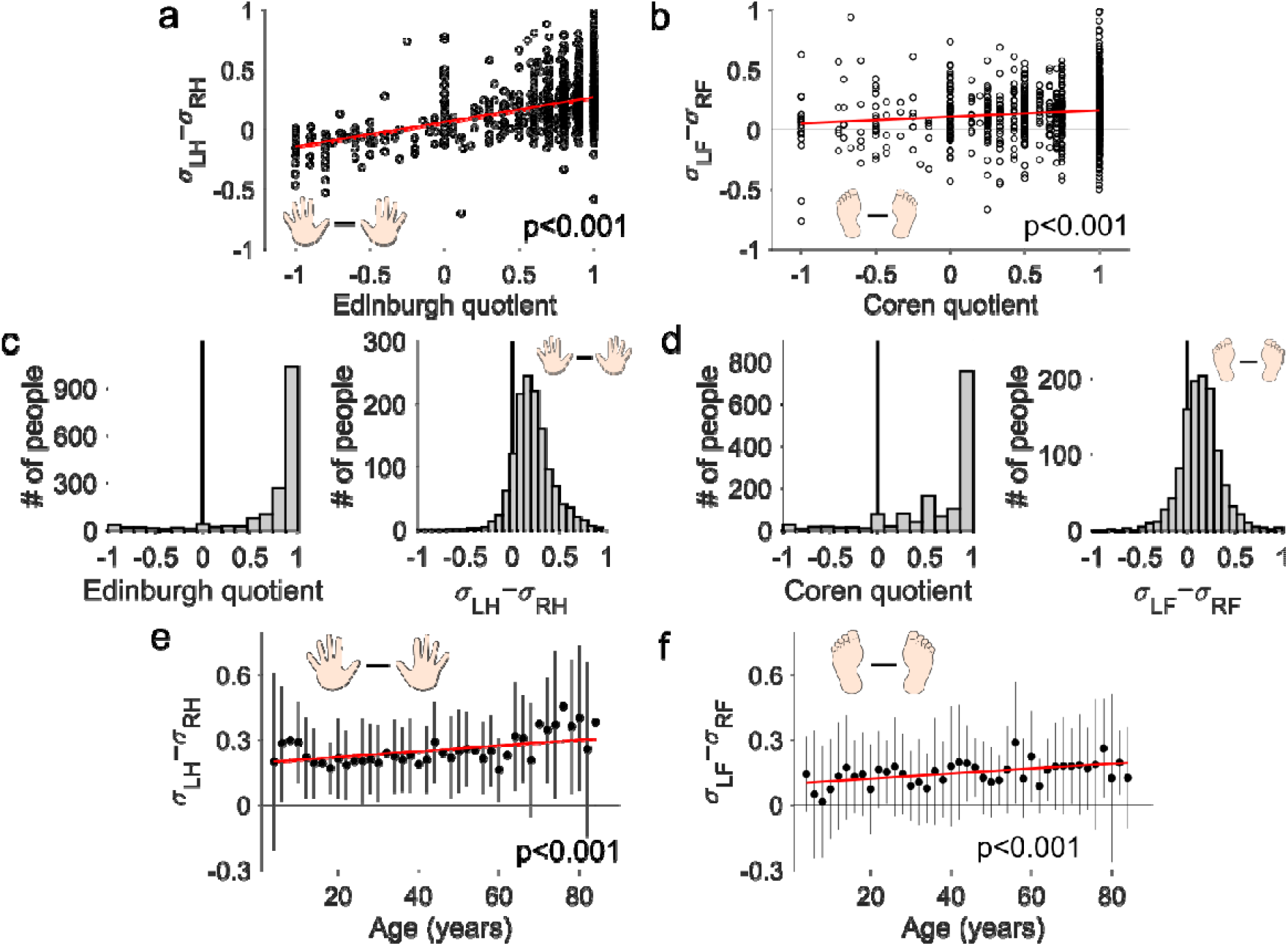
Left minus right acceleration variability of the hands and feet are related to handedness and footedness, respectively. (a) Left-hand minus right-hand acceleration variability was positively related to the Edinburgh quotient. (b) Left-foot minus right-foot acceleration variability was weakly related to the Coren quotient. (c) Histograms of and the Edinburgh quotient. The Edinburgh quotient shows a J-shaped distribution while shows a heavy-tailed distribution. (d) Histograms of and the Coren quotient. Coren quotient has a J-shaped distribution like the Edinburgh quotient, and is distributed more closely to a normal distribution. (e) Difference in left minus right-hand acceleration variability increased with age due to faster regression of the non-dominant hand. (f) The laterality of the feet also increased with age as the non-dominant foot regressed more than the dominant foot.

The distributions of *σ*_LH_ − *σ*_RH_, *σ*_LF_ − *σ*_RF_, and the Edinburgh and Coren quotients were examined by their histograms to determine how sensitive they were in measuring handedness and footedness (Figures 3c and 3d). The Edinburgh and Coren quotients showed a J-shaped distribution as they clustered at -1 and +1, while *σ*_LH_ − *σ*_RH_ and *σ*_LF_ − *σ*_RF_ were closer to a normal distribution. The clustering of the Edinburgh and Coren quotients is due to a ceiling effect where most participants answered that they used the right hand or foot for all tasks. The Edinburgh and Coren quotients are good at identifying whether someone is left- or right-handed and left- or right-footed, but they are limited at determining the degree of handedness and footedness.

We next looked at whether *σ*_LH_ − *σ*_RH_ and *σ*_LF_ − *σ*_RF_, which serve as markers of the laterality of the hands and feet, changed as a function of age (Figures 3e and 3f). Two linear mixed-effects models were fit to *σ*_LH_ − *σ*_RH_ and *σ*_LF_ − *σ*_RF_ separately. This analysis revealed that the slope was significantly positive for both *σ*_LH_ − *σ*_RH_ (slope=0.0013±0.0003, p<0.001) and for *σ*_LF_ − *σ*_RF_ (slope=0.001±0.0003, p<0.001). Thus, both *σ*_LH_ − *σ*_RH_ and *σ*_LF_ − *σ*_RF_ increased with age. This is due to the faster regression of the non-dominant hand and foot relative to the dominant limbs (Figures 2c and 2d).

Does practice change the acceleration variability of the hands? We examined the effect of prolonged practice on motor skill by focusing on dominant hand correction. In our study, 102 left-handed individuals reported that they had been forced to write with their right-hand as a child (out of which data from the feet was available from n=83). First, we looked at the acceleration variability of the left and right hands from three groups (Figure 4a): forced-handed individuals (n=102), strongly left-handed individuals with an Edinburgh quotient less than -0.5 (n=48), and strongly right-handed people with an Edinburgh quotient greater than +0.5 (n=1188). We carried out a two-way repeated-measures ANOVA with the hand’s acceleration variability as the dependent variable and the side (left-hand, right-hand) as a repeated categorical variable and the group (strongly left-handed, strongly right-handed, forced-handed) as a categorical variable. This analysis revealed a significant main effect of the side (F(1,1335)=36.3, p<0.001) and group variables (F(2, 1335)=8.6, p<0.001) as well as their interaction (F(2, 1335)=112.9, p<0.001) on the hand’s acceleration variability. Post-hoc multiple comparisons using Tukey’s HSD revealed that the left-hand acceleration variability *σ*_LH_ of strongly left-handed and forced-handed individuals was similar (p=0.08), and the right-hand’s acceleration variability *σ*_RH_ of strongly right-handed and forced-handed people was also comparable (p=0.10). The forced-handers’ *σ*_LH_ was significantly smaller than that of strongly right-handed people (p<0.001), and the forced-handers’ *σ*_RH_ was also smaller than that of strongly left-handed individuals (p<0.001). Forced-handed individuals had low acceleration variability in both their left and right hands, a pattern that was different from both strongly left-handed and right-handed individuals.

**Figure 4.**
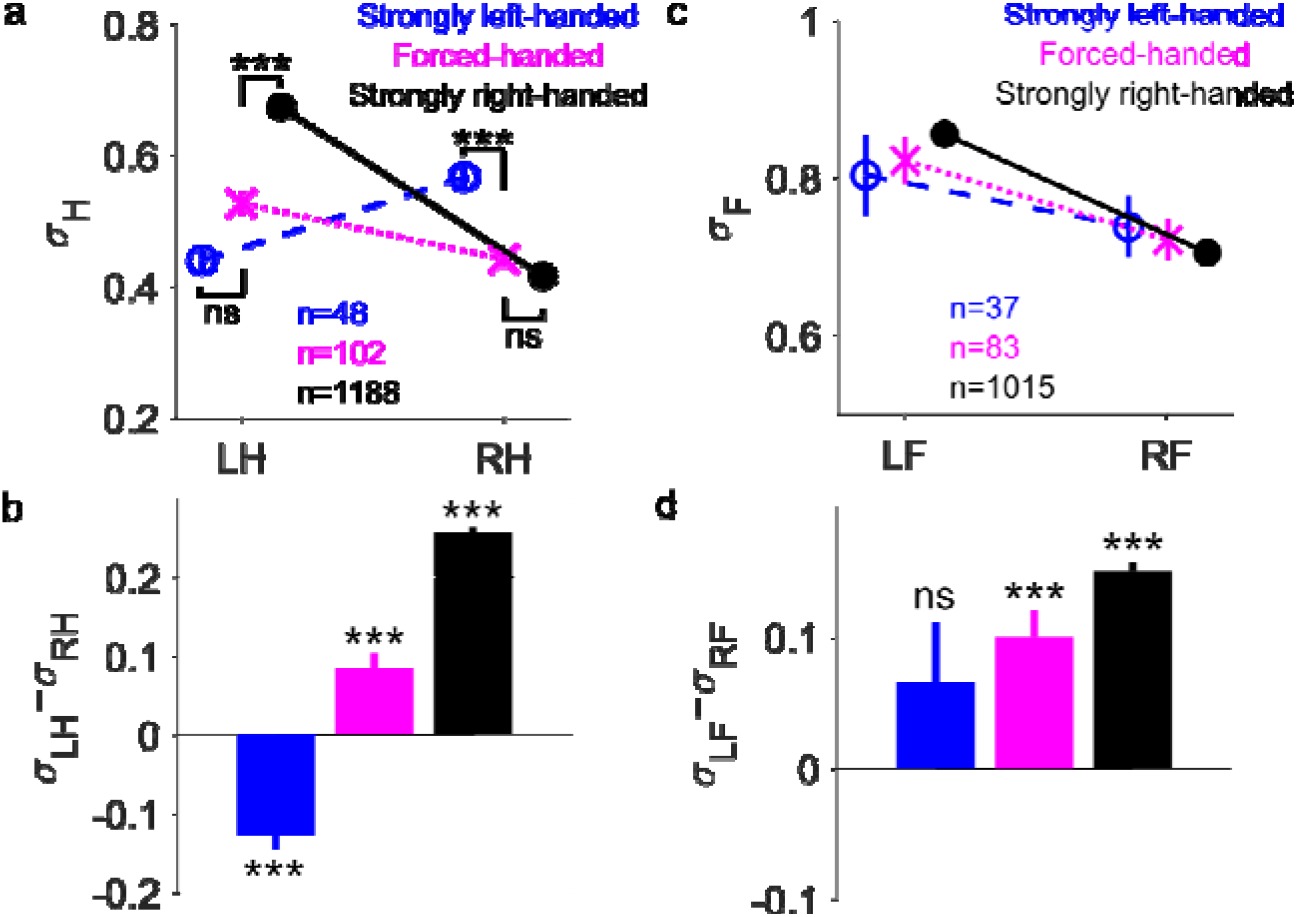
Forced use of the right-hand affects the relative acceleration variability between the left- and right-hands and the left- and right-feet. (a) Acceleration variability of the left and right-hands from strongly left-handed people (blue), strongly right-handed people (black), and forced-handed people (magenta). Two-way repeated measures ANOVA revealed that both the left- and right-hand acceleration variability of forced-handed individuals were not significantly different from the acceleration variability of the dominant hands of strongly left-handed and strongly right-handed people (*** signifies p<0.001). (b) Left-hand minus right-hand acceleration variability from strongly left-handed, strongly right-handed, and forced-handed individuals. One-sample t-tests with Holm-Bonferroni corrections revealed that of forced-handed people was significantly greater than zero, shifting closer to that of strongly right-handed individuals. (c) Acceleration variability of the left and right feet from strongly left-handed, strongly right-handed, and forced-handed individuals. The Edinburgh quotient was used to separate the groups. (d) Left-foot minus right-foot acceleration variability of strongly left-handed, strongly right-handed, and forced-handed individuals. One-sample t-tests revealed that forced-handed individuals’ was significantly greater than zero like strongly right-handed individuals.

Does forced correction affect the laterality of the hands? To answer this question, we examined the left-hand minus the right-hand’s acceleration variability *σ*_LH_ − *σ*_RH_ from strongly left-handed, strongly right-handed, and forced-handed individuals (Figure 4b). One-sample t-tests with a Holm-Bonferoni correction revealed that *σ*_LH_ − *σ*_RH_ of forced-handed individuals was significantly greater than zero (*t*(101)=4.53, p<0.001), which is closer to the *σ*_LH_ − *σ*_RH_ of strongly right-handed individuals rather than strongly left-handed ones. Thus, forced use of the right-hand reduced the right-hand’s acceleration variability without compromising the left-hand’s acceleration variability. The change in *σ*_LH_ − *σ*_RH_ suggests a shift towards right-handedness when right-hand usage is forced upon originally left-handed individuals, which shows that motor learning reduces the movement variability.

Previous research has shown how the right-handed participants show a preference in using the right-foot for manipulation like kicking a ball^19^ and that the right-foot of right-handers can tap faster than the left-foot^20^. This suggests a connection or transfer between hand and foot skill that may be genetic or learned through experience. If training or experience may affect hand and foot skill, could prolonged training of the right-hand transfer to the right-foot, thereby increasing the right-foot’s skill? To check whether forced correction of the hand affects the motor skill of the feet, we plotted the left-foot and the right-foot’s acceleration variability from strongly left-handed, strongly right-handed, and forced-handed individuals (Figure 4c). The groups were separated by handedness and not footedness. We carried out a two-way repeated-measures ANOVA with the foot’s acceleration variability as the dependent variable and the side (left-foot, right-foot) as a repeated categorical variable and the group (strongly left-handed, strongly right-handed, forced-handed) as a categorical variable. This revealed that the side (F(1,1132)=59.4, p<0.001) and the interaction between the side and the group (F(2,1132)=5.1, p=0.006) significantly affected the foot’s acceleration variability, but the main effect of the group was not significant (F(2,1132)=0.08, p=0.92). Post-hoc comparisons with Tukey’s HSD revealed no significant differences between the foot’s acceleration variability amongst any of the groups.

While the foot acceleration variability between groups was not significantly different, we checked whether the laterality of the feet or the left-minus right-foot’s acceleration variability *σ*_LF_ − *σ*_RF_ was different between the three groups (Figure 4d). One-sample t-tests with a Holm-Bonferroni correction showed that while *σ*_LF_ − *σ*_RF_ of strongly left-handed individuals was not different from zero (*t*(36)=1.43, p=0.15), it was significantly greater than zero in forced-handed (*t*(82)=5.14, p<0.001) and strongly right-handed individuals (*t*(1014)=23.5, p<0.001). While the skill of the left and right feet was not different in strongly left-handed individuals, both forced-handers and strongly right-handed individuals had superior right-foot motor skill. Thus, the laterality of the feet is significantly affected by prolonged training of the right-hand.

## Discussion

The aim of this study was to develop and validate an accessible assessment of motor skill based on the movements of the hands and feet. We used a smartphone to measure the acceleration of each limb as it traced a circle at a fixed rhythm and created an algorithm that quantified the variability of the three-axis acceleration trajectory. Our assessment was tested on 1675 participants of age three to eighty-eight years old to observe how the motor skill of the hands and feet changed with age and with prolonged training.

The acceleration variability of the hands and feet rapidly matured in the mid-teens and abruptly plateaued. This finding is consistent with other assessments of motor skill. The time taken to insert pegs into a series of holes declined and plateaued in the mid-teens^8^. The number of finger and foot taps also matures at adolescence^9^. The rapid growth and plateauing of motor skill in adolescence could be due to a maturation of the connections between the cerebellum and the cerebrum. The cerebellum and its connections to the thalamus and cerebrum play an important role in the execution of skillful actions^21^. These regions are involved in motor timing, which we have recently shown to be critical to determining the variability of movements^22,23^. Functional and diffusion MRI brain imaging has revealed that cerebrocerebellar connections are still immature in children and early adolescents relative to young adults^24^. The maturation of the cerebrocerebellar tract may be needed for the coordination of the hands and feet to reach their full potential.

The motor skill of both the hands and feet began to gradually regress after maturing in the mid-teens. This is also consistent with the literature as the ability to tap with the hand and foot declines with age^25^ and so does the performance when using a pencil to tap alternately between two lines^26^. One may believe that these changes in motor skill are due to weakening muscles, but peak muscular strength begins to decline from the twenties^27– 29^, which is significantly earlier than the regression observed in our study and in other studies^25,26^. The regression of motor skill observed in the elderly in our study may be attributed to the declining brain rather than muscle weakness. Ageing is known to reduce dopamine transmission, which reduces reaction times^30^ and leads to increased variability during standing^31,32^. The volume of white matter is also reduced in the elderly, leading to poorer fine motor performance^33^. Thus, changes in the brain’s structure and function may contribute to the regression in the motor skill of the hands and feet. Our study also revealed how the non-dominant hand and foot regressed faster than the dominant ones with aging. This is consistent with another study that found greater regression in the non-dominant hand when inserting pegs into holes^34^. Other studies have also shown that fine motor performance degrades faster in the non-dominant limb^35,36^. This would explain why laterality increased as a function of age in our data.

Left-handers who were forced to using their right hand from an early age had highly skilled left-and right-hands. The ability of forced-handers in using both the left and right hands skillfully may come from changes in both the brain’s function and its structure. A brain imaging study has shown that forced-handers display right-left asymmetry of central sulcus size, which is usually observed in right-handers^37^ and is an anatomical indicator of an individual’s handedness^38^. However, forced-handers continue to recruit higher-order premotor and parietal motor areas in the right hemisphere during handwriting and finger tapping^39,40^, which is more typical of left-handers. Thus, forced-handers possess a mixture of functional and structural features typical of both left-handers and right-handers. This is consistent with our results as forced-handers were essentially both left- and right-handed in the sense that the left- and right-hands were as skilled as the dominant hands of left-handers and right-handers. However, it remains unclear why forced use of the right-hand also affects the motor skill of the feet. One possibility is that increased use of the right-hand, e.g., to reach out and grab a cup, necessarily increases right-foot use as well to maintain balance. Another possibility is that the structural and functional changes during hand use are carried over to foot use as well, so changes in the laterality of the hands affects the feet as well.

The difference in the acceleration variability between the left and right limbs was related to both the handedness and the footedness of the participants, which was determined using the Edinburgh and Coren questionnaires. However, the distribution of the Edinburgh and Coren quotients and their tendency to cluster at the maximum values of -1 and +1 point to a limitation in the questionnaire approach at determining the degree of laterality of the hands and feet. Since most right-handed and right-footed individuals select the right-hand and right-foot for all items of the questionnaire, the Edinburgh and Coren quotients cannot discern the degree of right-handedness in strongly right-handed individuals. Other studies have raised a related issue when classifying individuals into left-handed, right-handed and mixed-handed groups^41,42^. Because these studies draw the line at different places and weigh the importance of questionnaire items differently, the same individual classified as mixed-handed in one study may be grouped into another category by others^43,44^. These problems highlight the importance of measuring the motor skill of the hands and feet to assess laterality rather than through questionnaires that are limited by a ceiling effect. A more graded measure of laterality would be useful in assessing the effects of training the non-dominant limb in sports and music training as pianists^45^ and drummers^46^ have been shown to exhibit less laterality or better non-dominant hand performance. Training of the weaker leg can also improve performance at soccer^47^. Our measure was sensitive enough to detect the improvement in non-dominant limb performance due to forced usage of the non-dominant hand. Hence, the newly proposed method could provide a convenient measure of changes in performance due to motor training.

## Methods

### Experiment protocol

The experiment was performed at either NTT Communication Science Laboratories or in Mizuno, both located in Japan. The study was approved by the NTT Communication Science Laboratories ethics committee (ID: R03-011) and the Mizuno ethics committee (ID: 20230217-64, 20230519-86, 20230818-117, 20240105-170, 20240308-199) with all participants providing written informed consent prior to participation. A guardian provided written informed consent on the behalf of underaged participants. A total of 1675 participants partook in this experiment.

When measuring hand dexterity, participants were instructed to hold a smartphone (Oppo A35, China) in either their left or right hand with the screen facing up at a comfortable position in front of them. Participants younger than 12 years of age used a mini smartphone (Melrose 2019, China). An A4-sized guide sheet was held in the other hand that had a dotted circle of diameter 10 cm printed on it. Participants traced the circle in time with a metronome that sounded from the smartphone at a speed of 2.5 Hz. They had to trace one circle per period for 15 seconds, after which the metronome stopped. They then switched hands and traced the circle with the other hand for 15 seconds. The order of measurement was chosen by the participant.

After both hands were measured, the smartphone was placed inside a running armband (Elecom eclear, Japan) that was strapped to the leg just above the ankle parallel to the sagittal plane. Participants placed both hands on a wall or solid object and stood straight. An A4-sized guide sheet printed with a 15 cm diameter circle was placed on the floor underneath the foot to be measured. Participants kept their leg straight and used their hip to trace the circle with their heel in time with the smartphone’s metronome set to 1.5 Hz for 15 seconds. Once one foot was measured, the smartphone was strapped to the other foot and the process was repeated. Like with the hands, the ordering of the measurement was decided by the participant.

The movement direction was clockwise for the right limb and anti-clockwise for the left limb. In the analysis, the first 1.5 seconds of data was discarded from as participants tended to begin moving sometime after the metronome started. The acceleration along the *x, y*, and *z* axes was collected for the entire duration of the movement. The readings from the accelerometer were captured at the maximum possible frequency, which was approximately 400 Hz using the Oppo A35 and 100 Hz using the smaller Melrose 2019, yielding roughly 6000 or 1500 data points per limb, respectively.

### Estimating movement variability from the acceleration trace

The movement variability was estimated from the three-axis acceleration trace through a processing pipeline. First, the offset along each axis was removed, and then the accelerometer readings were divided by its average Euclidean vector norm to resize the acceleration trace for comparison purposes. Since the movement was periodic, the acceleration trace could be separated into revolutions. To separate the data into revolutions, we used principal component analysis (PCA) to find the two principal components with the highest variance. The acceleration trace was then rotated in three dimensions so that the first two principal component vectors were along the *x* and *y* axes. One revolution was defined to be the time when the phase angle crossed 0° and the next time it did so. For 13.5 seconds worth of data, 31.2±0.1 revolutions for the hand and 18.1±0.1 for the foot were identified.

With the accelerometer data divided into revolutions, we compared the similarity of the acceleration trajectory between neighboring revolutions to avoid drifts from affecting the results. There are several ways of comparing the similarity of two trajectories. They can be broadly classified into metric measures like the Euclidean distance and the Fréchet distance^48^ and non-metric measures like dynamic time-warping^49^ and the longest common subsequence^50^. In our case, a metric measure is desirable as the measure should be zero if the trajectories are identical and it should obey the triangular inequality^51^. Furthermore, the measure should be time insensitive since the sampling rate of the two trajectories depends on what other tasks the smartphone is doing when recording the accelerometer’s data. For these reasons, we chose the Hausdorff distance measure^17^, which is commonly used in image analysis, to find the maximum Euclidean distance between the two furthest points on the two revolutions. These comparisons were made for all neighboring revolutions pairs, and the average Hausdorff distance yielded the acceleration variability for the left hand *σ*_LH_, right hand *σ*_RH_, left foot *σ*_LF_, and right foot *σ*_RF_ .

### Polynomial fitting of the hand and foot acceleration variability data

We conducted a series of linear-mixed affects analysis on the hand and foot acceleration data above the age of maturation to determine the order of the polynomial fit to use. We gradually added higher order terms to the model and used likelihood ratio tests to see if the term significantly improved the fit. This was carried out for the non-dominant and dominant hand and foot data separately (a total of four data sets). The likelihood ratio tests were only significant up to the second-order term for the hand and non-dominant foot data. For the dominant foot data, it was significant until the third-order term, but to allow comparisons between all data sets a second-order polynomial was used to fit all data above the age of maturation.

### Bootstrap resampling

The confidence intervals of the age of maturation *Y*_M_ was obtained by the bootstrap resampling method. Taking the hand data as an example, if *N* is the total number of participants, then the acceleration variability of the dominant and non-dominant hands from the same participant were sampled *N* times from the entire dataset with replacement. The value of *Y*_M_ that minimized the sum of squares distance between the data and the two polynomial fits (one linear fit before *Y*_M_ and a second-order fit after *Y*_M_) were obtained for each resampling set, and 1.96 times the standard deviation of the 10000 values of *Y*_M_ was used as the 95% confidence interval of *Y*_M_ . The same process was used separately to obtain the confidence intervals of *Y*_M_ for the hands and feet separately.

## Notes

### Competing Interest Statement

The authors have declared no competing interest.

